# Relationships between age, fMRI correlates of familiarity and familiarity-based memory performance under single and dual task conditions

**DOI:** 10.1101/2023.05.26.542526

**Authors:** Marianne de Chastelaine, Erin D Horne, Mingzhu Hou, Michael D Rugg

**Author notes:** Corresponding author: Marianne de Chastelaine, Center for Vital Longevity and School of Behavioral and Brain Sciences, University of Texas, Dallas, TX 75235, USA.

## Abstract

Using fMRI, we investigated the effects of age and divided attention on the neural correlates of familiarity and their relationship with memory performance. At study, word pairs were visually presented to young and older participants under the requirement to make a relational judgment on each pair. Participants were then scanned while undertaking an associative recognition test under single and dual (auditory tone detection) task conditions. The test items comprised studied, rearranged (words from different studied pairs) and new word pairs. fMRI familiarity effects were operationalized as greater activity elicited by studied pairs incorrectly identified as ‘rearranged’ than by correctly rejected new pairs. The reverse contrast was employed to identify ‘novelty’ effects. Behavioral familiarity estimates were equivalent across age groups and task conditions. Robust fMRI familiarity effects were identified in several regions, including medial and superior lateral parietal cortex, dorsal medial and left lateral prefrontal cortex, and bilateral caudate. fMRI novelty effects were identified in the anterior medial temporal lobe. Both familiarity and novelty effects were age-invariant and did not vary according to task condition. In addition, the familiarity effects correlated positively with a behavioral estimate of familiarity strength irrespective of age. These findings extend a previous report from our laboratory, and converge with prior behavioral reports, in demonstrating that the factors of age and divided attention have minimal impact on behavioral and neural estimates of familiarity.

## 1. Introduction

The impact of age on memory performance depends on whether the memory judgment requires recollection of contextual details about a study event or can be based on an acontextual sense of familiarity. Whereas age-related memory decline is prominent when performance depends on recollection (e.g., see Koen and Yonelinas, 2014; Old and Naveh-Benjamin, 2008, for reviews), the accuracy of familiarity-based memory judgments is markedly less sensitive to age (see Koen and Yonelinas, 2014; Yonelinas, 2002, for reviews). An influential proposal regarding the distinction between recollection and familiarity is that, while recollection depends on controlled processing, familiarity is relatively automatic (Jacoby, 1991). If this proposal is correct, it implies that recollection is more resource limited than familiarity. Since the availability of domain general processing resources declines in older age (Craik, 2020; Craik & Byrd, 1982), this may be one reason why recollection is more vulnerable than familiarity to increasing age.

Numerous behavioral studies have investigated resource limitations on memory performance using the divided attention (DA) paradigm, whereby memory performance is contrasted according to whether encoding or retrieval is undertaken without distraction or while also performing a secondary task. The effects of DA at retrieval on memory performance, the focus of the current study, are generally found to be more subtle than the effects of DA at encoding (e.g., Baddeley et al., 1984; Craik et al., 1996). Notably, studies examining the impact of DA at retrieval on estimates of familiarity and recollection in young adults have led to mixed results, but largely report minimal, or null, effects on both measures (Craik et al., 2018; Gruppuso et al., 1997; Knott and Dewhurst, 2009; Rosenstreich and Goshen-Gottstein, 2015; Skinner and Fernandes, 2008). Instead, DA at retrieval is more often reported to impact the secondary task, primarily reflected by increased RTs under dual task relative to single task conditions (e.g., Baddeley et al., 1984; Craik et al., 2018). These findings support the notion that memory retrieval may be in some way ‘protected’, such that processing resources are obligatorily allocated to retrieval processing and away from the secondary task (Craik et al., 1996).

Only a handful of studies have examined the effects of DA at retrieval on familiarity estimates (see above), and only one of these (Skinner and Fernandes, 2008) included groups of young and older adults. In this study, Skinner and Fernandes (2008) reported a detrimental impact of DA on familiarity estimates when the memory and secondary tasks shared similar stimulus materials (words), but no effect of DA on familiarity when task materials differed (words and digits, respectively). These differential effects of DA on familiarity estimates were reported to be age-invariant. Here, we employed fMRI and a DA manipulation at retrieval with groups of young and older adults to further evaluate the effects of resource limitations on behavioral estimates of familiarity, along with their neural correlates (i.e., fMRI ‘familiarity effects’), and any moderating effects of age. To our knowledge, this is the first study to describe the impact of DA at retrieval on fMRI familiarity effects in either young or older participants.

fMRI familiarity effects, often operationalized by the contrast between recognized but unrecollected items and unstudied items, typically take the form of enhanced activity for familiar items in a distributed set of brain regions that include the intra-parietal sulcus (IPS), precuneus, dorsal medial and anterolateral prefrontal cortex (dmPFC and alPFC respectively), and the caudate nucleus (de Chastelaine et al., 2017; Hou et al., 2021; Kim, 2010, 2013). Relatively greater familiarity is also associated with activity reductions, sometimes referred to as ‘novelty effects’, in other regions (e.g., Tulving and Kroll, 1995; see Kafkas and Montaldi, 2014, for review). Most investigations of such familiarity-related activity reductions have focused on the anterior medial temporal lobe (MTL), notably the perirhinal cortex (PRC) (e.g., Henson et al., 2003; Staresina et al., 2012; Wang et al., 2014; see Diana et al., 2007, for review of early studies) and anterior hippocampus (e.g., Daselaar et al., 2006; Staresina et al., 2012; see Kim, 2013, Nyberg, 2005, and Rugg et al., 2012, for reviews). Novelty effects, however, have also been identified in other regions, including mPFC, temporoparietal junction (TPJ), middle and posterior cingulate gyrus and insula (e.g., Angel et al., 2013; de Chastelaine et al., 2017; Hou et al., 2021).

Only a few studies have investigated the effects of age on fMRI correlates of familiarity-based memory judgments (Angel et al., 2013; Daselaar et al., 2006; de Chastelaine et al., 2017; Duarte et al., 2010; Hou et al., 2021; Wang and Giovanello, 2016). While some studies report null effects of age (Daselaar et al., 2006; Hou et al., 2021; Wang and Giovanello, 2016), others have reported familiarity-related enhancement of cortical activity in some regions to be either attenuated (Angel et al., 2013; de Chastelaine et al., 2017; Duarte et al., 2010) or exaggerated (Duarte et al., 2010) with increased age. Findings from fMRI studies that have examined the influence of age on novelty effects and novelty processing more generally indicate that both cortical and MTL novelty-related processing is largely preserved with increasing age (Angel et al., 2013; Daselaar et al., 2006; de Chastelaine et al., 2017; Duarte et al., 2010; Hou et al., 2021; Moriguchi et al., 2011; Wang and Giovanello, 2016; Wang et al., 2015; see Bowman and Dennis, 2015, for an exception), although age-related enhancements have been reported for novelty effects in the perirhinal cortex (Daselaar et al., 2006) and the hippocampus (Wang et al., 2015).

In a previous fMRI study (de Chastelaine et al., 2017), we employed an associative memory procedure to investigate the effects of age on behavioral and neural indices of familiarity in young, middle-aged and older participants. Consistent with the prior literature, behavioral estimates of familiarity-based discrimination were only modestly affected by age relative to recollection performance. fMRI familiarity effects were identified in several cortical regions previously identified as familiarity sensitive, and fMRI novelty effects were identified in, among other regions, bilateral anterior hippocampus and perirhinal cortex. Except for familiarity effects in one small region of the dmPFC, the effects were age-invariant and demonstrated age-invariant positive relationships with memory performance across participants, suggesting little influence of age on the processes underlying familiarity-based recognition.

Here, again employing an associative memory procedure, we follow up on these findings in young and older adults, but in the context of a DA manipulation at retrieval. As in our previous study (de Chastelaine et al., 2017), we expected to find minimal effects of age on both the behavioral and neural indices of familiarity-related processing when attention was undivided. In light of previous behavioral findings indicating minimal effects of DA at retrieval and, given the proposal that familiarity processing is minimally affected by resource depletion (Jacoby, 1991) and that, in any case, retrieval processing is ‘protected’ (Craik, 1996), we expected DA to have minimal impact on behavioral estimates of familiarity in either the young or older group. The key question was whether DA would impact fMRI familiarity or novelty effects in either age group. Even if there is little or no impact of DA on behavioral estimates of familiarity, it remains to be determined whether this extends to familiarity-related neural activity. For example, one possible scenario is that preserved memory performance under DA requires compensatory recruitment of neural resources, reflected perhaps by increased activity within previously identified familiarity-sensitive regions or the recruitment of additional regions. If this were the case, we might expect such compensatory activity to be greater for older than for young adults, given the decline in general processing resources that occurs with increasing age (Craik, 2020; Craik & Byrd, 1982).

## 2. Materials and methods

Data from this experiment were reported in two prior publications (Horne et al., 2021; Hou et al., 2022) that examined the effects of DA on fMRI monitoring- and recollection-related effects respectively. Further details of the methods can be found in those reports. The behavioral estimates of familiarity, and the familiarity- and novelty-related fMRI findings that we describe below have not been reported previously.

### 2.1. Participants

Twenty-four young (19-30 years) and 26 older (65-76 years) cognitively healthy adults participated in the current experiment. Data collected from 6 additional individuals (4 young and 2 older) were excluded because of insufficient trial numbers (i.e., less than 7 trials) for fMRI analysis in one of the critical conditions. Participants gave informed consent in accordance with the UT Dallas and UT Southwestern Institutional Review Boards. See Horne et al. (2021) for details of the participant inclusion and exclusion criteria. All participants completed a standard battery of neuropsychological tests on a day prior to that of the MRI session. See Horne et al. (2021) for details.

### 2.2. Materials

The experimental materials comprised 320 semantically unrelated word pairs that were randomly divided into five lists of 64 pairs. The five lists were rotated across participants such that each list provided items for each of three experimental word pair categories: intact, rearranged and new (see below). For each participant, word pairs from four of the lists were used to create a study list. Critical items for the test list included the 192 pairs that had been presented at study (intact pairs), 64 pairs of studied words that had been re-paired from study (rearranged pairs), and 64 pairs of unstudied words (new pairs). The test items were intermixed with 104 null trials. For both the study and test lists, the different categories of word pairs were pseudorandomized so that no more than three pairs from the same category occurred successively. Two buffer pairs were presented at the beginning and in the middle of the study and test lists.

A randomly determined sequence of low (400 Hz) and high (900 Hz) frequency auditory tones was presented concurrently with each test list. Stimulus onset asynchrony (SOA) for the tones ranged between 1000-3000 ms. To avoid the possibility of cross-modal perceptual interference, tone onsets did not occur during the display of a red fixation cross (presented 500ms prior to each word pair), or during the first 500 ms of the word pair presentation. ‘Target’ and ‘non-target’ tones (see below) were presented in a 30:70 ratio. Target tone frequency (high or low) was counterbalanced across participants.

### 2.3. Experimental procedure

Participants were given instructions and practice sessions for the study task, the memory test and tone detection task prior to scanning. Participants were therefore aware that their memory for the study items would be tested. As part of the practice session, participants performed the tone detection task in the absence of any test items. Accuracy and response time (RT) from this session were used to define baseline performance for the task. Tone detection performance, and the procedures employed to ensure that participants were able to comfortably perceive the tones in the scanner, are described in Horne et al. (2021).

The study phase was administered outside the scanner on a laptop computer and included two study blocks separated by a brief rest interval. The study task required participants to indicate with a button press which of the two objects denoted by the words was more likely to fit into the other. Following the study phase, participants were taken to the scanner and prepared for the test phase. The scanned test phase began approximately 25 minutes after completion of the study phase. The test phase included four consecutive blocks separated by short rest intervals. Test blocks alternated between single task (associative memory task only) and dual task (associative memory plus tone detection) conditions. In the single task condition, participants were instructed to ignore the tones and focus on the memory test. In the dual task condition, participants were asked to perform the memory and tone detection tasks concurrently and to give equal emphasis to each task. For the memory test, participants were instructed to press one of three keys, using the index, middle and ring fingers of one hand, to indicate whether a test pair was intact, rearranged or new. An ‘intact’ response was required when both words were recognized and there was a specific memory of the two words having been presented together at study.

A ‘rearranged’ response was required when both words were recognized from the study phase but there was no specific memory of the words having been paired together previously. Participants were instructed to make a ‘new’ response when neither or only one word was recognized. The tone detection task required participants to signal with a key press using the index finger of the other hand whenever a target tone occurred.

In both the study and test phases, word pairs were presented for a duration of 2000 ms, one word above and one below a central fixation character. The words were presented in white uppercase 30-point Helvetica font against a black background. Word pairs were preceded by a red fixation cross presented for 500 ms and followed by a white fixation cross for 1000 ms at study and for 2000 ms at test. Null trials during the scanned test phase consisted of the presentation of a white fixation cross against a black background for 4500 ms. A rest period of 30 s was included midway through each test block. For the test phase, hand assignments, response-finger mapping and the ordering of the task conditions were counterbalanced across participants.

### 2.4. Behavioral analysis strategy

For each task condition, we computed an estimate of familiarity-based discrimination, pF, derived from the independence assumption underlying dual-process theories of recognition memory (Yonelinas and Jacoby, 1995). When the independence assumption holds, it has been argued that pF is a relatively ‘process pure’ measure of familiarity strength. This measure was estimated as [(p rearranged response to intact items)/(1-p correct response to intact items)] – [(p rearranged response to new items)/(1 - p intact response to new items)] and was subjected to a 2 (task) × 2 (age group) ANOVA. RTs for intact items incorrectly identified as rearranged (‘associative misses’) and unstudied pairs correctly endorsed as new (‘correct rejections’ or ‘CRs’) were entered into a 2 (task) × 2 (item type) × 2 (age group) ANOVA.

### 2.5. fMRI acquisition

A Philips Achieva 3T MRI scanner (Philips Medical System, Andover, MA USA) equipped with a 32-channel head coil was used to acquire functional and anatomical images. A 3D MP-RAGE pulse sequence (FOV=256×256, voxel size 1×1×1 mm, 176 slices, sagittal acquisition) was employed for T1-weighted anatomical image acquisition. Functional scans were acquired with a T2*–weighted echo-planar imaging sequence (TR 2 s, TE 30 ms, flip angle 70°). Each functional volume comprised 33 axial slices (3 mm thickness, 1 mm inter-slice gap) with an in-plane resolution of 3×3 mm. Slices were oriented parallel to the AC-PC line, acquired in ascending order, and positioned for full coverage of the cerebrum and most of the cerebellum. Functional data were acquired with a sensitivity encoding (SENSE) reduction factor of 2. The first five volumes of each block were discarded to allow tissue magnetization to achieve a steady state.

### 2.6. fMRI preprocessing

fMRI data were preprocessed and analyzed using SPM12 (http://www.fil.ion.ucl.ac.uk). Functional images were motion and slice-time corrected, realigned, and spatially normalized using a sample-specific template based on the MNI reference brain (Cocosco et al. 1997). The normalized images were then smoothed using an 8 mm full-width half-maximum (FWHM) Gaussian kernel. The functional data from the different test blocks were concatenated using the spm_concatenate.m function before being entered into the first level GLMs (see below). T1 anatomical images were normalized with a procedure analogous to that applied to the functional images and were employed to define anterior hippocampal and perirhinal regions of interest (ROIs; see below), as well as to localize and depict the functional results.

### 2.7. fMRI analysis strategy

Critical item types employed in the analysis of the fMRI data were associative misses and CRs. In line with prior fMRI investigations contrasting activity elicited by familiar and novel items (e.g., de Chastelaine et al., 2017; Henson et al., 1999), fMRI familiarity effects were operationalized as greater BOLD signal for associative misses than for CRs. Associative misses are assumed to have been wrongly endorsed as rearranged on the basis of an above-criterion familiarity signal in the absence of a recollection signal or when recollection was too weak to support an intact judgment (see de Chastelaine et al. 2016). By contrast, CRs are assumed to be associated with a low (sub-criterial) familiarity signal and absent recollection. We employed the reverse contrast (CRs > associative misses) to operationalize novelty effects (see Introduction). Items that might also be considered appropriate for these contrasts are those that were correctly identified as rearranged (rearranged hits), as these can be identified on the basis of familiarity alone. As we previously argued, however (de Chastelaine et al., 2016; see also de Chastelaine et al., 2017), rearranged hits can also be supported by a recollection-based, ‘recall-to-reject’ strategy (Rotello and Heit, 2000; Yonelinas and Parks, 2007). As recall-to-reject has been reported to be employed more frequently in young than in older participants (Cohn et al., 2008), employment of rearranged hits in the familiarity or novelty contrasts could potentially introduce a confound between age and use of this strategy.

### 2.8. fMRI analysis

In the first stage of the fMRI analysis, separate GLMs were constructed for each participant. BOLD activity elicited by test pairs was modeled as a delta function and convolved with a canonical hemodynamic response function (HRF) and a second, ‘delayed’, HRF generated by shifting an orthogonalized canonical HRF one TR (2s) later in time (the results associated with this basis function are not discussed further as they did not add anything of theoretical interest to the results reported below). Critical item types (associative misses and CRs) were separately modeled, along with correctly identified intact items and rearranged hits, for the single and dual tasks. All other item types, the 30-s rest breaks, the six motion regressors, the four constants modeling the mean BOLD signal for each test block, and spike regressors modeling volumes with a transient displacement relative to the previous volume of > 1mm or > 1 degree in any direction were modeled as covariates of no interest. Voxel-wise parameter estimates obtained for each participant from these first level GLMs were employed in the analyses described below.

#### 2.8.1. ROI analyses

Analyses of cortical familiarity effects (operationalized as greater BOLD signal for associative misses than for CRs) were conducted on participant-specific parameter estimates for each critical item type derived from six *a priori* defined ROIs: left intraparietal sulcus, left precuneus, left lateral anterior PFC, left dorsal PFC and bilateral caudate (see Figure 1A and Table 1 for peak coordinates and corresponding Brodmann Areas of these ROIs). The ROIs were defined based on the analyses of an independent dataset described in a previous report of the effects of age on the neural correlates of familiarity processing (de Chastelaine et al., 2017). Data were extracted from all voxels contained within 5 mm spheres (3 mm spheres for left and right caudate) centered on the peak voxels of clusters showing fMRI familiarity effects within each of the six ROIs and averaged. The mean parameter estimates were analyzed with a 2 (task) × 2 (item type) × 6 (region) × 2 (age group) ANOVA. We also conducted multiple regression analyses to assess the strength of any relationships between familiarity effects (associative misses - CRs) and memory performance. The regression model included pF, our metric of familiarity strength (see above) as the dependent variable, and age group, the fMRI familiarity effect (collapsing the parameter estimates across regions and task conditions to minimize the number of analyses) and the age group × fMRI familiarity effect interaction term as predictor variables. As the interaction was consistently non-significant, this was dropped from the models reported below. The outcomes of the regression analyses (and for the analogous analyses of the MTL novelty effects described below) are reported as partial correlation coefficients indexing the strength of the relationship between the familiarity effect and pF.

**Figure 1.**
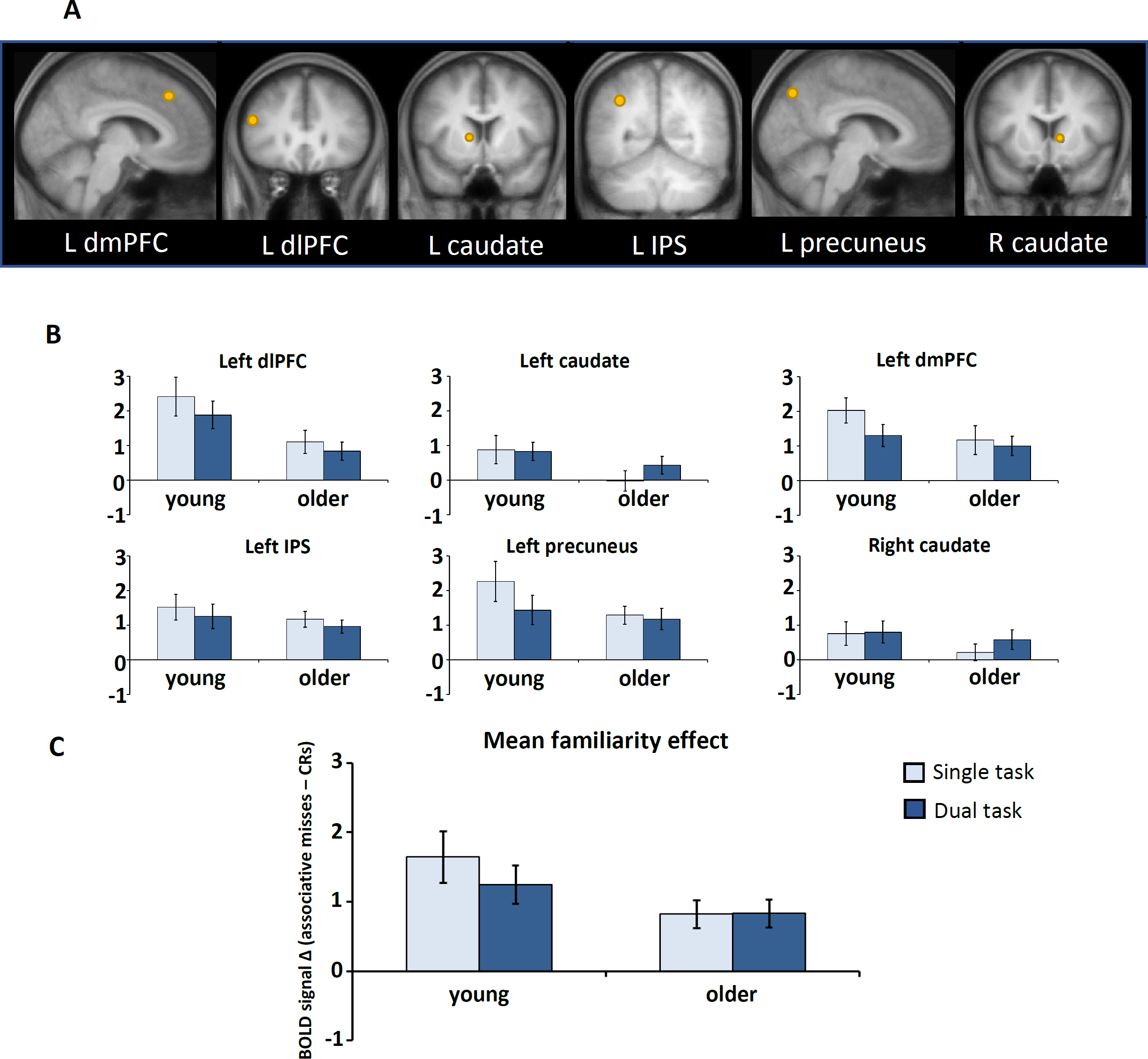
(A) Representative sagittal and coronal sections of the across-participants mean T1-weighted structural image depicting functional ROIs (orange spheres) centered on peak voxel coordinates from familiarity-sensitive cortical regions; (B) Mean familiarity effects (‘associative misses’ – ‘CRs’ parameter estimates) for each of the 6 *a priori*-determined ROIs; C) Mean familiarity effects averaged across the 6 ROIs. Error bars represent SEM.

**Table 1.** Peak coordinates and corresponding Brodmann Areas (BAs) of the 6 *a priori* defined familiarity-sensitive ROIs.

| Coordinates |  |  | Region | BA |
| --- | --- | --- | --- | --- |
| *x* | *y* | *z* |  |  |
| -6 | 29 | 44 | Left dorsal mPFC | 8 |
| -48 | 29 | 23 | Left anterior IPFC | 9 |
| -12 | 8 | 5 | Left caudate |  |
| -36 | -58 | 40 | Left IPS | 40 |
| -6 | -76 | 44 | Left Precuneus | 7 |
| 12 | 11 | 2 | Right caudate |  |

We examined fMRI novelty effects within anatomically defined MTL regions of *a priori* interest, namely, bilateral perirhinal cortex and anterior hippocampus. The ROIs were manually traced on the across-group average T1 anatomical template following the protocols specified by Insausti et al. (1998) (perirhinal cortex) and the EADC-ADNI Harmonized Protocol for Hippocampal Segmentation (Boccardi et al., 2015) (hippocampus). As proposed by Poppenk et al. (2013), the anterior hippocampus was defined as the portion of the hippocampus anterior to y = -21 in MNI space. The masks were constructed conservatively to ensure no overlap between the perirhinal and hippocampal regions. Thus, after reslicing and smoothing the ROIs to approximate the smoothness of the functional data, no voxels were shared between the perirhinal and hippocampal masks (see Figure 3A). Parameter estimates for associative misses and CRs were averaged across all voxels within the ROIs.

Data from bilateral hippocampus and perirhinal cortex were entered into a 2 (task: single, dual) × 2 (item type: associative misses, CRs) × 2 (hemisphere; left, right) × 2 (region: hippocampus, perirhinal cortex) × 2 (group: young, older) ANOVA to examine whether the effects in these MTL regions interacted with task condition or age group, and whether the profile of the effects differed according to region or hemisphere. Preliminary analyses revealed that the factor of hemisphere did not interact with any other factor and, therefore, for the purposes of the analyses reported here, the data were collapsed across this factor. As for the cortical familiarity effects, we also conducted multiple regression analyses to examine the relationships between novelty effects and memory performance separately for hippocampal and perirhinal ROIs, using the difference between the parameter estimates (CR – associative misses) as a predictor variable. To minimize the number of regression analyses conducted, we again averaged the parameter estimates across hemispheres and, additionally, across task conditions. The regression models included pF as the dependent variable, and age group, the fMRI novelty effect, and the age group × fMRI novelty effect interaction term as predictor variables. As for the familiarity-sensitive ROIs, the interaction terms were not significant and so were dropped from the models reported below.

Bayes factors (BF) were estimated to assess the strength of the evidence supporting any theoretically significant null findings that arose from the null hypothesis significance testing approach described above. We report BF inclusion (BF_incl_) values for the ANOVAs. For each of these estimates, values > 1 support the alternative hypothesis while values < 1 are considered to support the null hypothesis (Jeffreys, 1961).

#### 2.8.2 Whole brain analysis

To complement the ROI analyses, we conducted an exploratory whole brain analysis of the fMRI data, which is reported in Supplemental Materials (see Figure S1 and Table S1). For this analysis, participant-specific parameter estimates taken from the first level GLM for correctly identified intact items, associative misses, rearranged hits and CRs were brought forward to a 2 (task) × 4 (item type) × 2 (age group) mixed-design ANOVA model as implemented within SPM12. An initial 2 (task) × 2 (item type: associative misses; CRs) × 2 (age group) ANOVA contrast was run at the whole brain level to identify where fMRI effects differed according to task condition and age group. This analysis was subjected to a height threshold of p < .001 and a cluster extent threshold of p < .05 after FWE correction. Using the same thresholds, fMRI effects common to young and older participants were identified with the across-group main effects of familiarity (associative misses > CRs) and novelty (CRs > associative misses), exclusively masked with the two-sided (associative misses vs CRs) age group × fMRI interaction contrast, liberally thresholded at p < .05 uncorrected.

Analyses were conducted using SPM12 (http://www.fil.ion.ucl.ac.uk), SPSS 27.0 and JASP 0.14.0.0. The Greenhouse-Geisser correction was applied to ANOVA contrasts where appropriate. Apart from the whole brain analysis, significance levels for all tests were set at p < .05.

## 3. Results

### 3.1. Behavior

Summaries of the behavioral data are presented in Tables 2 and 3. Table 2 shows proportions of intact, rearranged and new pairs according to memory judgment, as well as a summary of the familiarity estimate, pF, for each task condition and age group. One sample t-tests indicated that pF was greater than chance in both young and older groups for each task condition (ts > 8.92, ps < .001, Cohen’s ds > 1.74). A 2 (task) × 2 (age group) ANOVA of pF failed to reveal any significant effects: age group [F(1,48) = 1.42, p = .239, partial η^2^ = 0.03, BF_incl_ = 0.54]; task condition [F(1,48) = 0.96, p = .333, partial η^2^ = 0.02, BF_incl_ = 0.32]; age group × task [F(1,48) = 0.00, p = .975, partial η^2^ = 0.00, BF_incl_ = 0.28].

**Table 2.** Mean proportions (SD) of intact, rearranged, and new test pairs given intact, rearranged, and new judgments, and pF in each age group and task condition. Correct responses are highlighted in bold. for each task condition and age group.

|  | Young |  | Older |  |
| --- | --- | --- | --- | --- |
|  | Single | Dual | Single | Dual |
| **Intact judgments** |  |  |  |  |
| Intact pairs | **0.68 (0.14)** | **0.66 (0.14)** | **0.57 (0.16)** | **0.54 (0.22)** |
| Rearranged pairs | 0.18 (0.12) | 0.19 (0.11) | 0.26 (0.18) | 0.31 (0.18) |
| New pairs | 0.04 (0.04) | 0.03 (0.05) | 0.09 (0.09) | 0.12 (0.11) |
| **Rearranged judgments** |  |  |  |  |
| Intact pairs | 0.19 (0.08) | 0.21 (0.11) | 0.28 (0.14) | 0.28 (0.15) |
| Rearranged pairs | **0.56 (0.14)** | **0.57 (0.13)** | **0.45 (0.17)** | **0.38 (0.15)** |
| New pairs | 0.22 (0.15) | 0.24 (0.14) | 0.28 (0.12) | 0.26 (0.14) |
| **New judgments** |  |  |  |  |
| Intact pairs | 0.13 (0.10) | 0.13 (0.07) | 0.15 (0.06) | 0.18 (0.10) |
| Rearranged pairs | 0.26 (0.11) | 0.24 (0.09) | 0.29 (0.14) | 0.30 (0.14) |
| New pairs | **0.74 (0.16)** | **0.72 (0.17)** | **0.63 (0.15)** | **0.62 (0.17)** |
| **pF** | 0.38 (0.16) | 0.36 (0.18) | 0.34 (0.17) | 0.31 (0.18) |

Table 3 shows mean response times (RTs) for the item types of interest – associative misses and CRs. A 2 (task condition) × 2 (item type) × 2 (age group) ANOVA revealed main effects of task condition [F(1,48) = 21.04, p < .001, partial η^2^ = 0.31] and item type [F(1,48) = 72.85, p < .001, partial η^2^ = 0.60], indicating, across the two age groups, slower responses in the single, relative to the dual, task condition and slower responses for associative misses than for CRs. There was no significant effect of age group [F(1,48) = 0.26, p .614, partial η^2^ = 0.01, BF_incl_ = 0.61], and there were no significant interactions between age group and task condition [F(1,48) = 0.14, p = .713 partial η^2^ = 0.00, BF_incl_ = 0.26], age group and item type [F(1,48) = 0.18, p = .671, partial η^2^ = 0.00, BF_incl_ = 0.50] or age group, task condition and item type [F(1,48) = 0.194, p = .662, partial η^2^ = 0.00, BF_incl_ = 0.16].

**Table 3.** Mean RT (SD) of associative misses and CRs in each age group and task condition.

|  | Young |  | Older |  |
| --- | --- | --- | --- | --- |
|  | Single | Dual | Single | Dual |
| **Associative misses** | 2172 (457) | 2043 (413) | 2094 (341) | 1996 (333) |
| **CRs** | 1890 (412) | 1798 (408) | 1851 (328) | 1762 (309) |

### 3.2. fMRI results

#### 3.2.1. Familiarity effects

Figures 1B and 1C illustrate the familiarity effects (i.e., difference in BOLD activity between associative misses and CRs) for each of the 6 functional ROIs and the across-ROI mean, according to age group and task condition. Table S2 (Supplemental Materials) summarizes the results of a 6 (region) × 2 (task) × 2 (item type) × 2 (age group) mixed effects ANOVA of the data extracted from each ROI. As the table indicates, the ANOVA revealed a significant main effect of item type [F(1,48) = 47.05, p < .001, partial η^2^ = 0.50], indicating, across the ROIs, greater activity for associative misses than for CRs (i.e., a significant familiarity effect). The familiarity effect did not differ significantly as a function of age group [F(1,48) = 1.40, p > .3, partial η2 = 0.02, BF_incl_ = 0.37] although there was a main effect of ROI [F(3.28,157.41) = 31.98, p < .001, partial η2 = 0.40]. The main effects of item type and ROI were qualified by interactions between these two factors [F(3.45,165.59) = 12.48, p < .001, partial η2 = 0.21] and between them and task [F(4.14,198.47) = 2.51, p < .05, partial η2 = 0.05].

To unpack this three-way interaction, we ran additional 2 (task condition) × 2 (item type) repeated measures ANOVAs for each ROI. The ANOVAs revealed a main effect of item type for each ROI (all ps < .01) which did not interact with task condition in any region (all ps > .1, all BF_incl_ values < 1.09). Thus, the origin of the interaction is obscure and is not discussed further.

As indicated in Table S3 (Supplementary Materials), partial correlation analysis (controlling for age group) revealed an age-invariant positive relationship between fMRI familiarity effects (mean across the 6 ROIs) and pF, averaged across task conditions (r = 0.32, p < .025). The partial plot illustrating this relationship is depicted in Figure 2.

**Figure 2.**
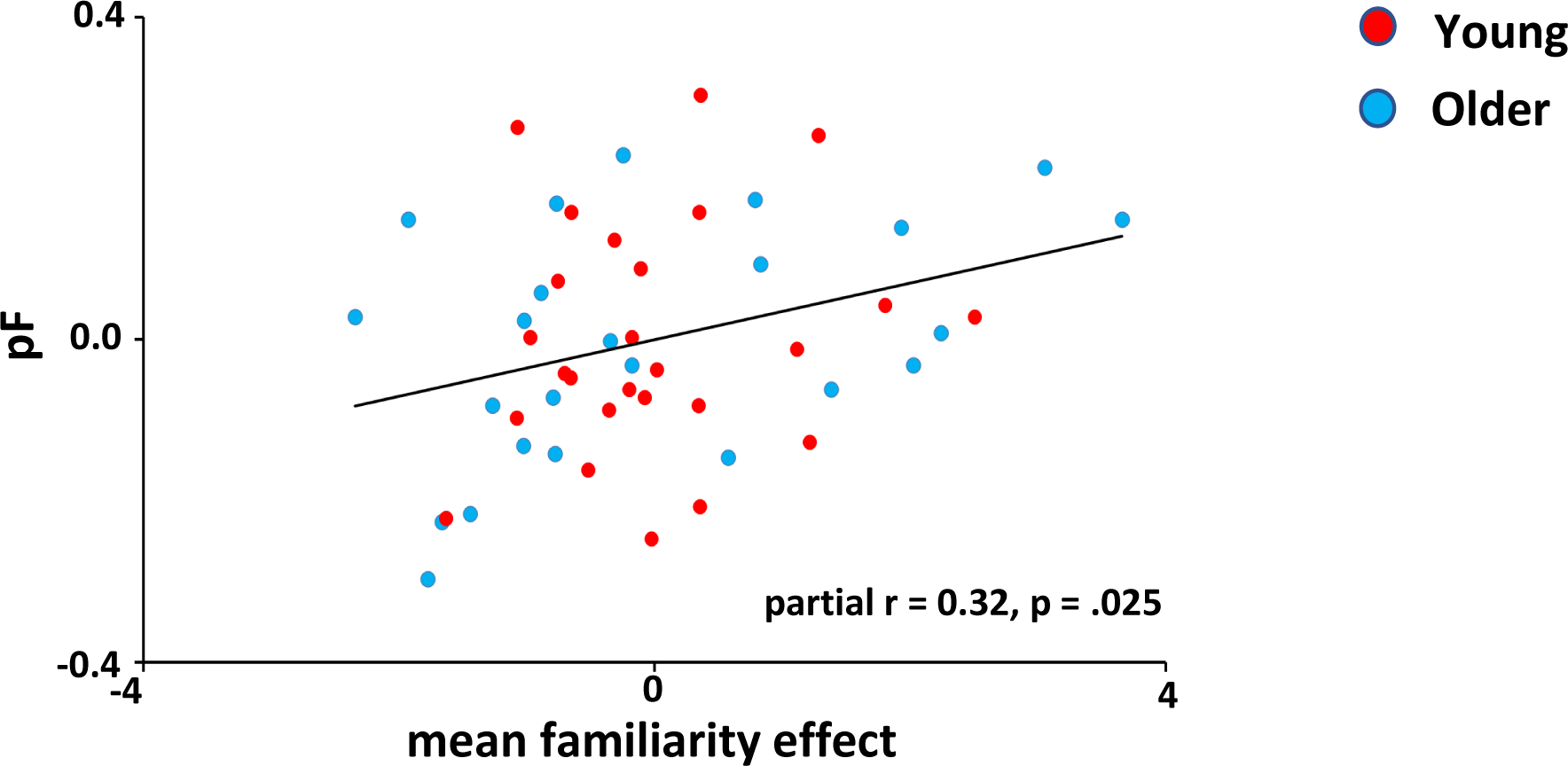
Partial plot showing the relationship across participants between mean familiarity effects and pF, averaged across task conditions, controlling for age group.

#### 3.2.2. Novelty effects

Figures 3B and 3C illustrate the novelty effects (i.e., difference in BOLD activity elicited by CRs and associative misses) for the hippocampal and perirhinal ROIs, and mean novelty effects across the 2 ROIs, respectively, according to age group and task. As is evident from Table S4 (Supplementary Materials), a 2 (age group) × 2 (item type) × 2 (task condition) × 2 (region) mixed design ANOVA gave rise to a reliable main effect of item type [F(1,48) = 5.00, p < .05, partial η^2^ = 0.09], indicating greater activity for CRs than for associative misses (i.e., a significant novelty effect). The novelty effect did not differ significantly as a function of age group [F(1,48) = 0.06, p > .08, partial η^2^ = 0.00] or task condition [F(1,48) = 1.76, p > .19, partial η^2^ = 0.04]. However, there were significant interactions between age group and task condition [F(1,48) = 4.66, p < .05, partial η2 = 0.09], age group and region [F(1,48) = 7.20, p < .001, partial η2 = 0.13], and age group, task condition and region [F(1,48) = 4.70, p < .05, partial η2 = 0.09]. To unpack the three-way interaction, we ran follow-up ANOVAs for each MTL region. For the hippocampus, the ANOVA revealed a cross-over age group × task condition interaction [F(1,48) = 10.03, p < .005, partial η2 = 0.17], such that mean hippocampal BOLD activity for the young group was significantly greater in the single task relative to the dual task condition [t(23) = 2.30, p < .05, Cohen’s d = 0.46] while, in the older group, this task condition difference was reversed [t(25) = 2.35, p < .05, Cohen’s d = 0.46] (see Figure 4A). The ANOVA for the perirhinal data failed to reveal a main effect of task condition [F(1,48) = 0.23, p > .6, partial η2 = 0.01, BF_incl_ = 0.23] or an interaction between age group and task condition [F(1,48) = 0.01, p > .9, partial η2 = 0.00, BF_incl_ = 0.29]. Across tasks, however, there was a main effect of age group, indicating greater item-related activity in the perirhinal cortex for the older group than for the young group [F(1,48) = 8.14, p < .01, partial η2 = 0.15] (see Figure 4B).

**Figure 3.**
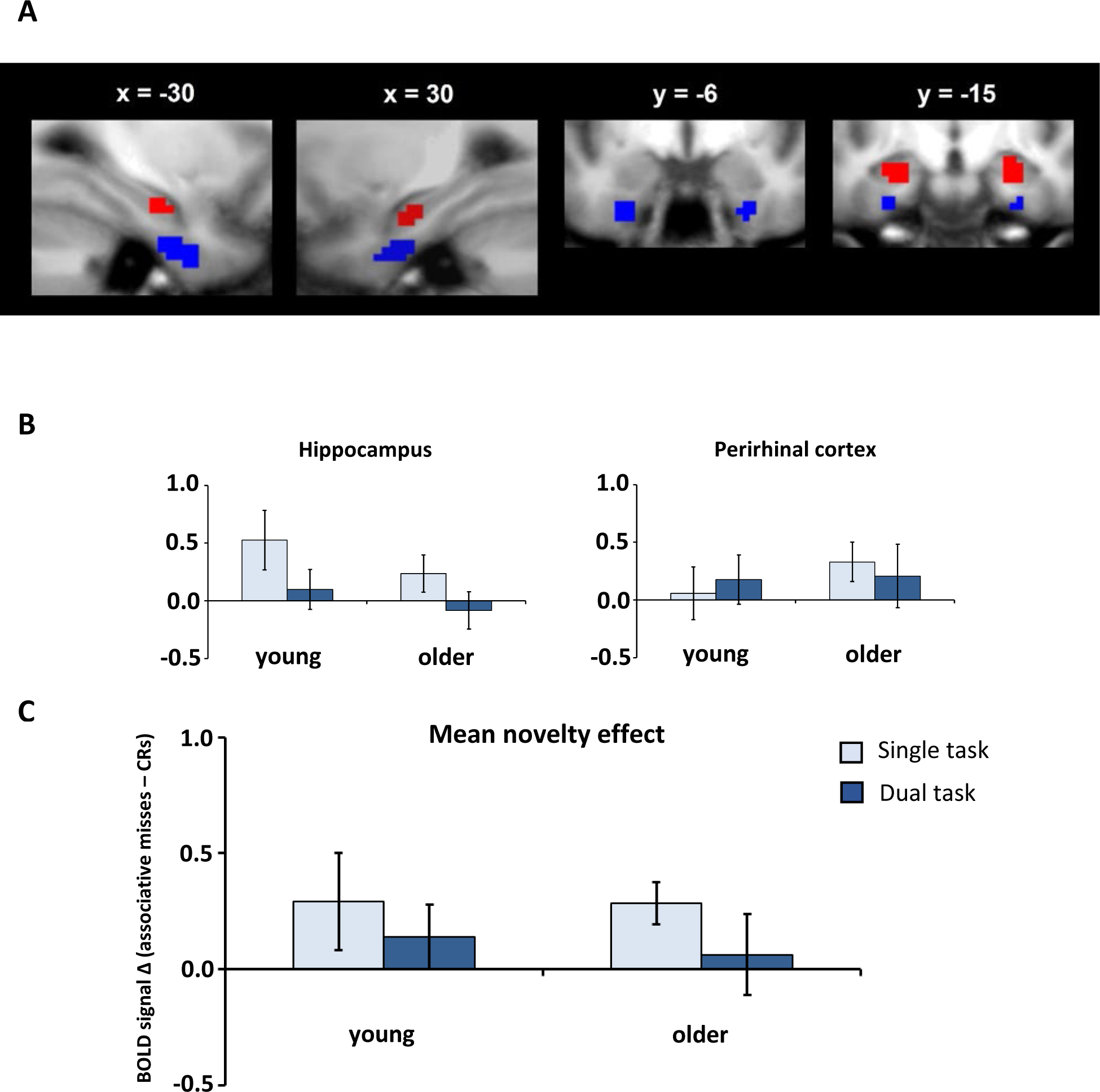
(A) Representative sagittal and coronal sections of the across-participants mean T1-weighted structural image showing manual tracings of the anterior hippocampus (red ROIs) and perirhinal cortex (blue ROIs); (B) Mean novelty effects (‘CRs’ – ‘associative misses’ parameter estimates) for the hippocampal and perirhinal ROIs; C) Mean novelty effects averaged across the 2 MTL ROIs. Error bars represent SEM.

**Figure 4.**
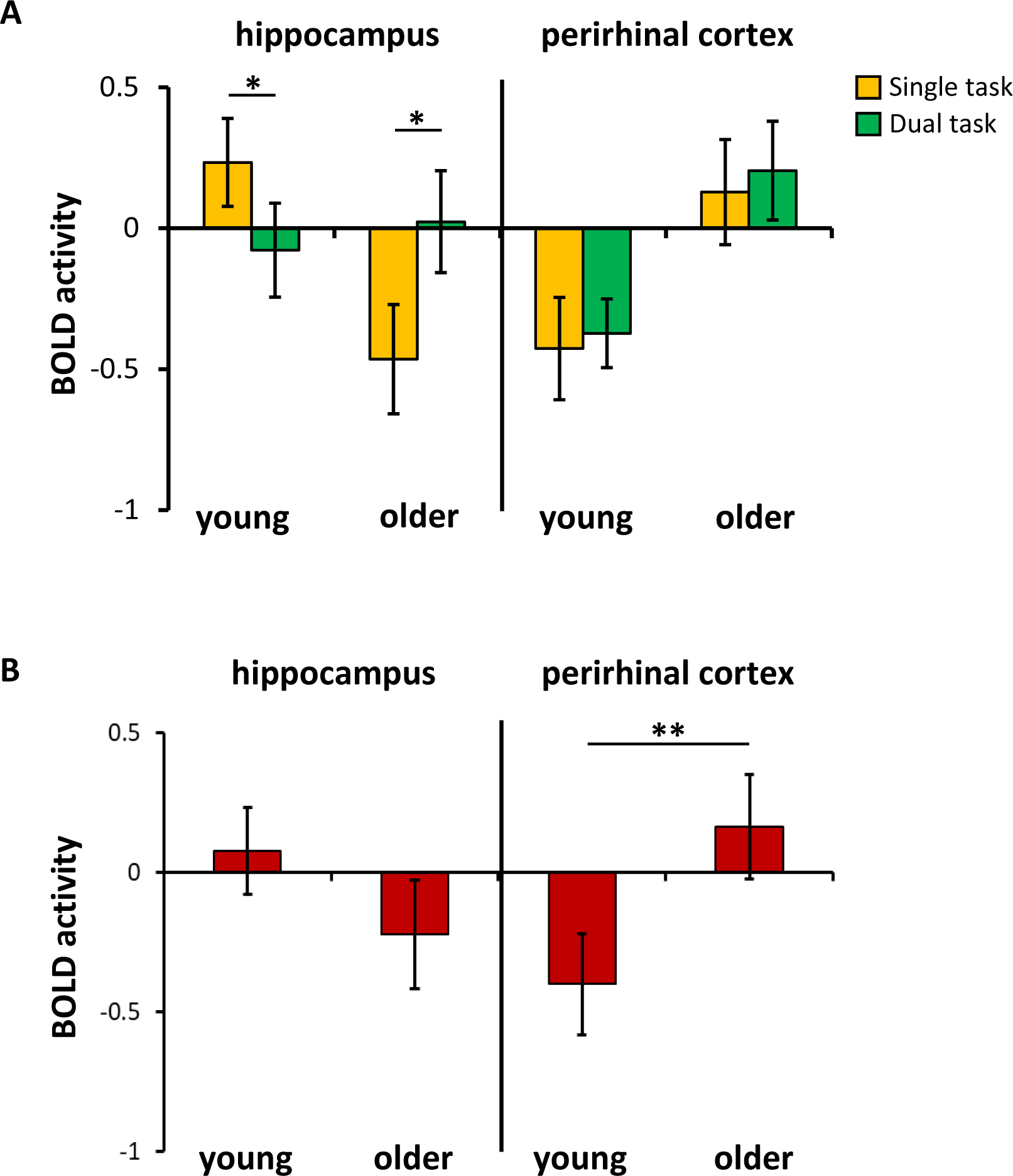
(A) Significant age group × task effect for hippocampal BOLD activity; (B) Significant effect of age group for perirhinal BOLD activity. BOLD activity is in arbitrary units, error bars represent SEM. * p < .05; ** p < .01.

Partial correlation analyses (controlling for age group) failed to reveal any significant relationships between hippocampal or perirhinal novelty effects and pF averaged across task conditions (ps > .2) (see Tables S5 and S6 in Supplementary Materials for a summary of these results).

## 4. Discussion

The current study examined the effects of DA at retrieval on familiarity-driven recognition memory and its fMRI correlates in young and older adults. Consistent with most previous findings (see Introduction and below), DA failed to impact behavioral estimates of familiarity in either age group. Across the two groups, robust cortical/striatal familiarity and MTL novelty effects were identified; these too were unaffected by the DA manipulation. However, mean (item-related) hippocampal BOLD activity for the older group was significantly greater in the dual task relative to the single task condition whereas this difference was reversed for the young group. Across tasks, item-related activity in the perirhinal cortex was greater for the older group than for the young group. Below, we expand on these findings and the other findings reported in the current experiment.

### 4.1. Behavior

In keeping with the majority of previous findings (see Introduction), familiarity strength (estimated by pF) did not significantly differ between the two age groups. Moreover, DA had no discernible impact on pF in either group. This null finding across the two age groups is consistent with the findings reported in a handful of previous studies of young adults only (Craik et al., 2018; Gruppuso et al., 1997; Knott and Dewhurst, 2009; Rosenstreich and Goshen-Gottstein, 2015), and in the one previous study to include both young and older adults (Skinner and Fernandes, 2008). Together, these findings indicate that processes supporting familiarity are minimally affected by resource depletion arising from DA at test and are consistent with the proposal that retrieval processing is in some way ‘protected’ in the face of the processing demands of a secondary task (Craik, 1996) (see Introduction).

### 4.2. fMRI findings

#### 4.2.1. Familiarity effects

fMRI familiarity effects were evident within each of the six *a priori* selected ROIs in both the young and older age groups. Crucially, the effects did not significantly differ between the two task conditions in either age group. Alongside the behavioral findings, these null results suggest that the processes supporting familiarity-based recognition judgments were minimally impacted by DA regardless of age. There was also little evidence of an influence of age on the familiarity effects. This latter finding is largely in agreement with our (de Chastelaine et al., 2017) and others’ (e.g., Angel et al., 2013; Daselaar et al., 2006; Duarte et al., 2010; Hou et al., 2021; Wang and Giovanello, 2016) prior reports that familiarity effects in most familiarity-sensitive regions are insensitive to age (see Introduction for details). The current null findings from the main ROI analysis are consistent with the outcome of the exploratory whole brain analysis, which also failed to identify an effect of DA or age on familiarity-related activity.

As in our prior report (de Chastelaine et al., 2017), and consistent with others’ previous findings (Daselaar, 2006; Hou et al., 2021; Johnson et al., 2013; Montaldi et al., 2006; Scimeca et al., 2016; Yonelinas et al., 2005), we identified an age-invariant positive association between the magnitude of familiarity effects and familiarity strength (pF) across participants. This finding adds to the evidence that fMRI familiarity effects track familiarity strength and do so in an age-invariant manner.

#### 4.2.2. Novelty effects

fMRI novelty effects, which were detected in the MTL ROIs, did not significantly vary by task condition in either group. Therefore, analogously with the behavioral familiarity estimates and fMRI familiarity effects, these null findings indicate that the processes reflected by MTL novelty effects are minimally affected by DA. The magnitude of the novelty effects also did not significantly differ between the young and older groups, consistent with our own (de Chastelaine et al., 2017) and others’ (Angel et al., 2013; Daselaar et al., 2006; Duarte et al., 2010; Hou et al., 2021; Moriguchi et al., 2011; Wang and Giovanello, 2016; Wang et al., 2015) previous reports that novelty-related processing in the MTL is largely preserved with increasing age.

Interpretations of novelty effects in the MTL have sometimes emphasized differential roles for the perirhinal cortex, thought to support familiarity-based recognition memory (e.g., Diana et al., 2007; Eichenbaum et al., 2007), and the hippocampus, proposed to support the encoding of novel items (e.g., Johnson et al., 2008; Köhler et al., 2005; Nyberg, 2005; Stark and Okado, 2003). However, it has also been suggested that the perirhinal cortex and hippocampus work synergistically to both detect and encode novel items, even within the context of a memory retrieval task (e.g., Fernandez & Tendolkar, 2006). As was noted in the Introduction, the effects of DA at encoding tend to be substantially larger than the effects of DA at retrieval on a wide number of memory measures, including estimates of familiarity. It has been suggested that relative to undivided attention at study, DA is likely to impair elaborative encoding, as indicated by reduced memory performance at test (Craik et al., 2018). The limited behavioral evidence suggests that, in the context of a retrieval task, DA impacts subsequent memory for the novel test items (Dudukovic and colleagues, 2009). Given the proposed role of the hippocampus and perirhinal cortex in the encoding of novel items during memory retrieval, it is therefore perhaps surprising that DA failed to impact MTL novelty effects in the present study. It will be important for future fMRI studies to examine the relationship between MTL novelty effects and subsequent memory performance for novel items in the context of a DA manipulation similar to that implemented here.

In contrast to the relatively robust relationship between MTL novelty effects and familiarity strength reported in our previous experiment (de Chastelaine et al., 2017), we were unable to identify any significant relationships between hippocampal or perirhinal novelty effects (identified by the ROI analysis) and familiarity estimates in the current experiment. Also, unlike in our previous study, exploratory whole brain analysis failed to identify *any* region that exhibited a novelty effect (or an interaction of an effect with task or age). It is possible, given our small sample size, that the current experiment merely lacked sufficient power to detect novelty effects given the conservative statistical thresholds necessitated by whole brain analysis. However, this seems unlikely given the robustness of the familiarity effects identified by the present whole brain analysis. An alternative possibility is that the weak novelty effects identified here reflect the relatively heavy task demands imposed by the experimental procedure. Notably, even in the single task condition, when attention was ostensibly undivided, participants nonetheless had to engage in the retrieval task while ignoring potentially distracting tones. While speculative, it is conceivable that novelty effects are reliant on attentional mechanisms that may not have been sufficiently available due to attentional capture by auditory tones, leading to weakened novelty effects and a breakdown in the relationship between MTL novelty effects and memory performance across task conditions.

#### 4.2.3. Item-related BOLD activity

Item-related BOLD responses (i.e., responses elicited by all classes of test item relative to baseline) in the hippocampus demonstrated a cross-over age group by task interaction, such that mean hippocampal BOLD activity in the young group was significantly greater in the single task relative to the dual task condition while, in the older group, mean hippocampal activity was greater in the dual task condition. Additionally, item-related activity in the perirhinal cortex was greater for older relative to young participants regardless of task condition. Age differences in item-related activity have previously been reported in mPFC (Hou et al, 2022), in mPFC and the hippocampus (Wang et al., 2016), and, in one study, across much of the ‘core recollection network’ (Hou et al., 2021). In each case, item-related activity was greater in older participants. The functional significance of age-related enhancements of item-related BOLD activity is uncertain, but one possibility is that it reflects upregulation of neural activity in response to increased cognitive challenge (cf. Cabeza et al., 2018). The present findings of greater perirhinal activity in older relative to young participants, and of greater hippocampal BOLD activity in the dual task compared to the single task in older adults, are arguably consistent with this proposal, but the finding of greater item activity in the single than in the dual task condition in the young participants is not. It will be important for future studies to further examine the functional significance of task-related modulations of item-related activity in young and older adults.

#### 4.2.4. Limitations

The present study has several limitations. First, although we did not identify any effects of age, the cross-sectional design of the study does not allow definitive conclusions to be drawn about the impact of *aging* on behavioral or neural correlates of familiarity (e.g., Raz and Lindenberger, 2011, Rugg, 2016). Second, given the modest sample sizes, caution is required in accepting null findings – in particular, limitations in sample size might have restricted the ability to detect subtle, but theoretically interesting, DA or age effects on brain-behavior relationships. Third, there are systematic age differences in the transfer function mediating between neural activity and the fMRI BOLD response (e.g., Liu et al., 2013; Lu et al., 2011; Tsvetanov et al., 2015). Since we did not control for this potential confound, its influence on the item-related activity findings we report cannot be ruled out. However, it seems unlikely that a generic age difference in the BOLD transfer function could fully explain our results – notably, the cross-over interaction between age and task effects in hippocampal item-related activity cannot be explained in these terms.

## 5. Conclusions

To our knowledge, this is the first study to examine the impact of DA on fMRI familiarity and novelty effects in either young or older adults. Across age groups, DA failed to influence either behavioral familiarity estimates or fMRI familiarity or novelty effects. The results support previous behavioral findings indicating that DA during a memory test has minimal impact on behavioral estimates of familiarity. The current study also extends these results by demonstrating that, regardless of age, neural correlates of familiarity processing are also unaffected by DA.

## Acknowledgements

## Acknowledgements

This work was supported by the National Institute on Aging [grant numbers R21AG054197, RF1AG039103].

## Declarations of Interest

None

## Notes

### Competing Interest Statement

The authors have declared no competing interest.

## References

Angel, L., Bastin, C., Genon, S., Balteau, E., Phillips, C., Luxen, A., Maquet, P., Salmon, E., Collette, F., 2013. Differential effects of aging on the neural correlates of recollection and familiarity. Cortex 49(6):1585–1597.

Baddeley, A., Lewis, V., Eldridge, M., & Thomson, N., 1984. Attention and retrieval from long-term memory. J. Exp. Psychol. Gen. 113, 518–540.

Boccardi, M., Bocchetta, M., Apostolova, L.G., Barnes, J., Bartzokis, G., Corbetta, G., DeCarli, C., Firbank, M., Ganzola, R., Gerritsen, L., Henneman, W., 2015. Delphi definition of the EADC-ADNI Harmonized Protocol for hippocampal segmentation on magnetic resonance. Alzheimers Dement. 11(2):126–38.

Bowman, C.R., Dennis, N.A., 2015. Age differences in the neural correlates of novelty processing: The effects of item-relatedness. Brain Res. 1612:2–15.

Cabeza, R., Albert, M., Belleville, S., Craik, F.I., Duarte, A., Grady, C.L., … & Rajah, M.N., 2018. Maintenance, reserve and compensation: the cognitive neuroscience of healthy ageing. Nature reviews. Neuroscience 19, 701–710.

Cocosco, C.A., Kollokian, V., Kwan, R.K.S., Pike, G.B., Evans, A.C., 1997. Brain web: Online interface to a 3D MRI simulated brain database. Neuroimage 5:425.

Cohn, M., Emrich, S.M., Moscovitch, M., 2008. Age-related deficits in associative memory: the influence of impaired strategic retrieval. Psychol. Aging 23(1):93–103.

Craik, F.I., & Byrd, M., 1982. Aging and cognitive deficits: The role of attentional resources. In Craik, F.I., Trehub, S.E. (Eds.), Aging and cognitive processes. Plenum Press. New York, pp. 191–211.

Craik, F.I., 2020. Remembering: An activity of mind and brain. Annu. Rev. Psychol. 71, 1–24.

Craik, F.I., Eftekhari, E., & Binns, M.A., 2018. Effects of divided attention at encoding and retrieval: Further data. Mem. Cogn. 46, 1263–1277.

Craik, F.I., Govoni, R., Naveh-Benjamin, M., & Anderson, N.D., 1996. The effects of divided attention on encoding and retrieval processes in human memory. J. Exp. Psychol. Gen. 125, 159–180.

Daselaar, S.M., Fleck, M.S., Dobbins, I.G., Madden, D.J., Cabeza, R., 2006. Effects of healthy aging on hippocampal and rhinal memory functions: an event-related fMRI study. Cereb Cortex 16(12):1771–1782.

de Chastelaine, M., Mattson, J.T., Wang, T.H., Donley, B.E., & Rugg, M.D., 2016. The neural correlates of recollection and retrieval monitoring: Relationships with age and recollection performance. Neuroimage 138, 164–175.

de Chastelaine, M., Mattson, J.T., Wang, T.H., Donley, B.E., & Rugg, M.D., 2017. Independent contributions of fMRI familiarity and novelty effects to recognition memory and their stability across the adult lifespan. NeuroImage 156, 340–351.

Diana, R.A., Yonelinas, A.P., Ranganath, C., 2007. Imaging recollection and familiarity in the medial temporal lobe: a three-component model. Trends Cogn Sci. 11(9):379–386.

Duarte, A., Graham, K.S., Henson R.N., 2010. Age-related changes in neural activity associated with familiarity, recollection and false recognition. Neurobiol. Aging 31(10):1814–1830.

Dudukovic, N.M., DuBrow, S., & Wagner, A.D., 2009. Attention during memory retrieval enhances future remembering. Memory & cognition 37(7), 953–961.

Eichenbaum, H, Yonelinas, A.P., Ranganath, C., 2007. The medial temporal lobe and recognition memory. Annu Rev Neurosci. 30:123–152.

Fernández, G., & Tendolkar, I., 2006. The rhinal cortex: ‘gatekeeper’ of the declarative memory system. Trends in cognitive sciences 10(8), 358–362.

Gruppuso, V., Lindsay, D.S., & Kelley, C.M., 1997. The process-dissociation procedure and similarity: Defining and estimating recollection and familiarity in recognition memory. Journal of Experimental Psychology: Learning, Memory, and Cognition 23(2), 259.

Henson, R.N., Rugg, M.D., Shallice, T, Josephs, O, Dolan, R.J., 1999. Recollection and familiarity in recognition memory: an event-related functional magnetic resonance imaging study. J Neurosci. 19(10):3962–3972.

Henson, R.N., Cansino, S., Herron, J.E., Robb, W.G., Rugg, M.D., 2003. A familiarity signal in human anterior medial temporal cortex? Hippocampus 13(2):301–304.

Horne, E.D., de Chastelaine, M., & Rugg, M.D., 2021. Neural correlates of post-retrieval monitoring in older adults are preserved under divided attention, but are decoupled from memory performance. Neurobiol. Aging 97, 106–119.

Hou, M., Horne, E.D., de Chastelaine, M., & Rugg, M.D., 2022. Divided attention at retrieval does not influence neural correlates of recollection in young or older adults. NeuroImage 250, 118918.

Hou, M., Wang, T.H., & Rugg, M.D., 2021. The effects of age on neural correlates of recognition memory: An fMRI study. Brain Cogn. 153, 105785.

Insausti, R., Juottonen, K., Soininen, H., Insausti, A.M., Partanen, K., Vainio, P., Laakso, M.P., Pitkänen, A., 1998. MR volumetric analysis of the human entorhinal, perirhinal, and temporopolar cortices. Am J Neuroradiol. 19(4):659–671.

Jacoby, L.L., 1991. A process dissociation framework: Separating automatic from intentional uses of memory. Journal of memory and language 30(5), 513–541.

Jeffreys, H., 1961. Theory of probability (3rd ed.). Oxford: Oxford University Press, Clarendon Press.

Johnson, J.D., Muftuler, L.T., Rugg, M.D., 2008. Multiple repetitions reveal functionally and anatomically distinct patterns of hippocampal activity during continuous recognition memory. Hippocampus 18(10):975–980.

Johnson, J.D., Suzuki, M., Rugg, M.D., 2013 Recollection, familiarity, and content-sensitivity in lateral parietal cortex: a high-resolution fMRI study. Front Hum Neurosci. 7(219):1–15.

Kafkas, A., Montaldi, D., 2014. Two Separate, But Interacting, Neural Systems for Familiarity and Novelty Detection: A Dual-Route Mechanism. Hippocampus 24:516–527.

Kim, H., 2010. Dissociating the roles of the default-mode, dorsal, and ventral networks in episodic memory retrieval. Neuroimage 50(4):1648–1657.

Kim, H., 2013. Differential neural activity in the recognition of old versus new events: An Activation Likelihood Estimation Meta-Analysis. Hum Brain Mapp. 34(4):814–836.

Knott, L.M., & Dewhurst, S.A., 2009. Investigating the attentional demands of recognition memory: Manipulating depth of encoding at study and level of attention at test. European Journal of Cognitive Psychology 21(7), 1045–1071.

Koen, J.D., Yonelinas, A.P., 2014. The effects of healthy aging, amnestic mild cognitive impairment, and Alzheimer’s disease on recollection and familiarity: a meta-analytic review. Neuropsychol. Rev. 24(3):332–354.

Köhler, S., Danckert, S., Gati, J.S., Menon, R.S., 2005. Novelty responses to relational and nonrelational information in the hippocampus and the parahippocampal region: a comparison based on event related fMRI. Hippocampus 15(6):763–774.

Liu, P., Hebrank, A.C., Rodrigue, K.M., Kennedy, K.M., Park, D.C., Lu, H., 2013. Age-related differences in memory-encoding fMRI responses after accounting for decline in vascular reactivity. Neuroimage 78:415–425.

Lu, H., Xu, F., Rodrigue, K.M., Kennedy, K.M., Cheng, Y., Flicker, B., Hebrank, A.C., Uh, J., Park, D.C., 2011. Alterations in cerebral metabolic rate and blood supply across the adult lifespan. Cereb. Cortex 21:1426–1434.

Montaldi, D., Spencer, T.J., Roberts, N., Mayes, A.R., 2006. The neural system that mediates familiarity memory. Hippocampus 16(5):504–520.

Moriguchi, Y., Negreira, A., Weierich, M., Dautoff, R., Dickerson, B.C., Wright, C.I., Barrett, L.F., 2011. Differential hemodynamic response in affective circuitry with aging: an FMRI study of novelty, valence, and arousal. J Cogn Neurosci. 23(5):1027–1041.

Nyberg, L., 2005. Any novelty in hippocampal formation and memory? Curr. Opin. Neurol. 18(4):424– 428.

Old, S.R., Naveh-Benjamin, M., 2008. Differential effects of age on item and associative measures of memory: a meta-analysis. Psychol Aging 23(1):104–118.

Poppenk, J., Evensmoen, H.R., Moscovitch, M., Nadel, L., 2013. Long-axis specialization of the human hippocampus. Trends Cogn Sci. 17(5):230–240.

Raz, N., & Lindenburger, U., 2011. Only time will tell: cross-sectional studies offer no solution to the age-brain-cognition triangle: comment on Salthouse, 2011. Psychol Bull 37:790-795.

Rosenstreich, E., & Goshen-Gottstein, Y., 2015. Recollection-based retrieval Is influenced by contextual variation at encoding but not at retrieval. PLoS One, 10(7), e0130403.

Rotello, C.M., Heit, E., 2000. Associative recognition: A case of recall-to-reject processing. Mem. Cognit. 28(6):907–922.

Rugg, M.D., 2016. Interpreting age-related differences in memory-related neural activity. In: Cabeza, R., Nyberg, L., Park, D.C., editors. Cognitive Neuroscience of Aging: Linking Cognitive and Cerebral Aging. 4th. Oxford University Press.

Rugg, M.D., Vilberg, K.L., Mattson, J.T., Yu, S.S., Johnson, J.D., Suzuki, M., 2012. Item memory, context memory and the hippocampus: fMRI evidence. Neuropsychologia 50(13):3070–3079.

Scimeca, J.M., Katzman, P.L., Badre, D., 2016. Striatal prediction errors support dynamic control of declarative memory decisions. Nat. Commun. 7(13061):1–15.

Skinner, E.I., & Fernandes, M.A., 2008. Interfering with remembering and knowing: Effects of divided attention at retrieval. Acta Psychologica 127(2), 211–221.

Staresina, B.P., Fell, J., Do Lam, A.T., Axmacher, N., Henson, R.N., 2012. Memory signals are temporally dissociated in and across human hippocampus and perirhinal cortex. Nat. Neurosci. 15(8):1167– 1173.

Stark, C.E., Okado, Y., 2003. Making memories without trying: medial temporal lobe activity associated with incidental memory formation during recognition. J. Neurosci. 23(17):6748–6753.

Tsvetanov, K.A., Henson, R.N., Tyler, L.K., Davis, S.W., Shafto, M.A., Taylor, J.R., Williams, N., Cam-Can Rowe, J.B., 2015. The effect of ageing on fMRI: Correction for the confounding effects of vascular reactivity evaluated by joint fMRI and MEG in 335 adults. Hum. Brain Mapp. 36:2248–2269.

Tulving, E., & Kroll, N., 1995. Novelty assessment in the brain and long-term memory encoding. Psychonomic bulletin & review 2(3), 387–390.

Wang, W.C., Dew, I.T., Cabeza, R., 2015. Age-related differences in medial temporal lobe involvement during conceptual fluency. Brain Res. 1612:48–58.

Wang, T.H., Johnson, J.D., de Chastelaine, M., Donley, B.E., Rugg, M.D., 2016. The effects of age on the neural correlates of recollection success, recollection-related cortical reinstatement, and post-retrieval monitoring. Cereb. Cortex 26:1698–1714.

Wang, W.C., Ranganath, C., Yonelinas, A.P., 2014. Activity reductions in perirhinal cortex predict conceptual priming and familiarity-based recognition. Neuropsychologia 52:19–26.

Wang, W.C., Giovanello, K.S., 2016. The role of medial temporal lobe regions in incidental and intentional retrieval of item and relational information in aging. Hippocampus 26(6):693–699.

Yonelinas, A.P., 2002. The nature of recollection and familiarity: A review of 30 years of research. J. Mem. Lang. 46(3):441–517.

Yonelinas, A.P., Otten, L.J., Shaw, K.N., Rugg, M.D., 2005. Separating the brain regions involved in recollection and familiarity in recognition memory. J. Neurosci. 25(11):3002–3008.

Yonelinas, A.P., Parks, C.M., 2007. Receiver operating characteristics (ROCs) in recognition memory: a review. Psychol. Bull. 133(5):800–832.

