## Supplemental data for "Relationships between age, fMRI correlates of familiarity and familiarity-based memory performance under single and dual task conditions"

*Whole brain analysis*

The initial 2 (task) x 2 (item type) x 2 (age group) ANOVA contrast at the whole brain level failed to identify any above threshold voxels where familiarity or novelty effects differed according to task condition or age group. The contrast also failed to identify any above threshold clusters exhibiting novelty effects across age groups. However, robust familiarity effects common to the two groups were identified in medial and lateral parietal cortex, dorsal medial and left lateral PFC, left caudate and right insula (see Figure S1 illustrating these effects and Table S1 for MNI coordinates of the peak voxels).

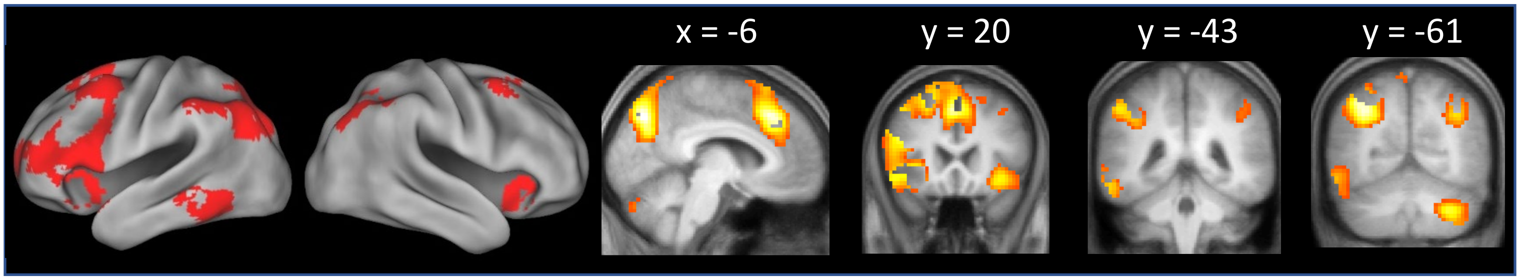


Figure S1. Clusters demonstrating familiarity effects common to the 2 age groups. The effects are superimposed on the bilateral surfaces of a standardized brain (PALS-B12) atlas using Caret 5, and on coronal sections of the across-group mean T1-weighted structural image.

Table S1. Peak voxels of the across-group main effect of familiarity, exclusively masked by the 2-sided age group by item type contrast.

| **Coordinates** | | | **Peak *Z*** | **No. of above-threshold voxels** | **Region** |
|---|---|---|---|---|---|
| ***x*** | ***y*** | ***z*** | | | |
| -3 | 26 | 44 | Infinite | 2916 | Left dorsal medial prefrontal cortex |
| -60 | -43 | -13 | 5.28 | 319 | Left inferior temporal gyrus |
| -6 | -76 | 44 | Infinite | 1716 | Left precuneus |
| 36 | 20 | -10 | 5.93 | 193 | Right insula |
| 36 | -61 | 38 | 5.55 | 403 | Right intraparietal sulcus |
| 30 | -67 | -37 | 7.58 | 634 | Right cerebellum |

Table S2. Summary of age group x item type x task condition x region ANOVA results for parameter estimates extracted from the 6 familiarity-sensitive cortical ROIs. Significant effects are highlighted with bold font. $BF_{incl}$ values are provided for non-significant results.

| Source | df | df error | F | p value | partial η2 | $BF_{incl}$ |
|---|---|---|---|---|---|---|
| age group | 1 | 48 | 0.43 | .515 | 0.01 | 0.37 |
| **item type** | **1** | **48** | **47.05** | **< .001** | **0.50** | |
| task | 1 | 48 | 0.56 | .458 | 0.01 | 0.22 |
| **region** | **3.28** | **157.41** | **31.98** | **< .001** | **0.40** | |
| age group x item type | 1 | 48 | 3.52 | .067 | 0.07 | 0.94 |
| age group x task | 1 | 48 | 3.89 | .054 | 0.08 | 0.77 |
| age group x region | 3.28 | 157.41 | 2.48 | .058 | 0.05 | 2.12 |
| item type x task | 1 | 48 | 1.04 | .314 | 0.02 | 0.35 |
| **item type x region** | **3.45** | **165.59** | **12.48** | **< .001** | **0.21** | |
| task x region | 3.39 | 162.66 | 2.18 | .084 | 0.04 | 0.05 |
| age group x item type x task | 1 | 48 | 1.17 | .286 | 0.02 | 0.01 |
| age group x item type x region | 3.45 | 165.59 | 1.31 | .270 | 0.03 | 0.00 |
| age group x task x region | 3.39 | 162.66 | 1.31 | .271 | 0.03 | 0.15 |
| **item type x task x region** | **4.14** | **198.47** | **2.51** | **.042** | **0.05** | |
| age group x item type x task x region | 4.14 | 198.47 | 0.35 | .850 | 0.01 | 0.00 |

Table S3. Results of the across-group regression models examining the relationships between mean familiarity effects (across the 6 familiarity-sensitive ROIs) and pF across task conditions: model includes age group as a covariate; b: unstandardized coefficient; SEb: standard error of the unstandardized coefficient; β: standardized coefficient.

| Model | b | SEb | β | partial r | p value |
|---|---|---|---|---|---|
| (constant) | 0.313 | 0.038 | | | |
| age group | -0.025 | 0.041 | -0.084 | -0.087 | .554 |
| familiarity effect | 0.040 | 0.017 | 0.327 | 0.320 | .025 |

Table S4. Summary of age group x item type x task condition x region ANOVA results for parameter estimates extracted from anterior hippocampal and perirhinal cortex ROIs. Significant effects are highlighted with bold font. $BF_{incl}$ values are provided for non-significant results.

| Source | F(1,48) | p value | partial n2 | $BF_{incl}$ |
|---|---|---|---|---|
| age group | 1.06 | .308 | 0.02 | 0.32 |
| **item type** | **5.00** | **.030** | **0.09** | |
| task | 0.65 | .424 | 0.01 | 0.46 |
| region | 0.08 | .779 | 0.00 | 0.35 |
| age group x item type | 0.06 | .810 | 0.00 | 0.27 |
| **age group x task** | **4.66** | **.036** | **0.09** | |
| **age group x region** | **7.20** | **.010** | **0.13** | |
| item type x task | 1.76 | .191 | 0.04 | 0.54 |
| item type x region | 0.00 | .988 | 0.00 | 0.20 |
| task x region | 0.02 | .899 | 0.00 | 0.27 |
| age group x item type x task | 0.06 | .813 | 0.00 | 0.14 |
| age group x item type x region | 1.98 | .166 | 0.04 | 0.50 |
| **age group x task x region** | **4.70** | **.035** | **0.09** | |
| item type x task x region | 1.95 | .169 | 0.04 | 0.45 |
| age group x item type x task x region | 0.43 | .514 | 0.01 | 0.53 |

Table S5. Results of the across-group regression models examining the relationships between hippocampal novelty effects and pF across task conditions: model includes age group as a covariate; b: unstandardized coefficient; SEb: standard error of the unstandardized coefficient; β: standardized coefficient.

| Model | b | SEb | β | partial r | p value |
|---|---|---|---|---|---|
| (constant) | 0.375 | 0.032 | | | |
| age group | -0.052 | 0.043 | -0.179 | -0.176 | .226 |
| hippocampal novelty effect | -0.011 | 0.032 | -0.053 | -0.053 | .720 |

Table S6. Results of the across-group regression models examining the relationships between perirhinal novelty effects and pF across task conditions: model includes age group as a covariate; b: unstandardized coefficient; SEb: standard error of the unstandardized coefficient; β: standardized coefficient.

| Model | b | SEb | β | partial r | p value |
|---|---|---|---|---|---|
| (constant) | 0.368 | 0.030 | | | |
| age group | -0.054 | 0.042 | -0.184 | -0.186 | .201 |
| perirhinal novelty effect | 0.029 | 0.024 | 0.171 | 0.173 | .236 |
